# Synonymous SNPs of viral genes facilitate virus to escape host antiviral RNAi immunity

**DOI:** 10.1101/496992

**Authors:** Yuechao Sun, Yu Zhang, Xiaobo Zhang

## Abstract

Synonymous single nucleotide polymorphisms (SNPs) are involved in codon usage preference or mRNA splicing. Up to date, however, the role of synonymous SNPs in immunity remains unclear. To address this issue, the SNPs of white spot syndrome virus (WSSV) were characterized in shrimp in the present study. Our results indicated that there existed synonymous SNPs in the mRNAs of wsv151 and wsv226, two viral genes of WSSV. In the presence of SNP siRNA, wild-type siRNA, wild-type mRNA and SNP mRNA of wsv151 or wsv226, RNAi was significantly suppressed, showing that the synonymous SNPs of wsv151 and wsv226 played negative roles in host siRNA pathway due to mismatch of siRNA with its target. In insect cells, the mismatch, caused by synonymous SNPs of wsv151 or wsv226, between siRNA and its target inhibited the host RNAi. Furthermore, the data revealed that the co-injection of SNP siRNA and wild-type siRNA of wsv151 or wsv226 into WSSV-infected shrimp led to a significant increase of WSSV copies compared with that of SNP siRNA alone or wild-type siRNA alone, indicating that the synonymous SNPs of viral genes could be a strategy of virus escaping host siRNA pathway in shrimp *in vivo*. Therefore, our study provided novel insights into the underlying mechanism of virus escaping host antiviral RNAi immunity by synonymous SNPs of viral genes.

**Author Summary:** Our results indicated that there existed synonymous SNPs in the mRNAs of wsv151 and wsv226, two viral genes of WSSV. In the presence of SNP siRNA, wild-type siRNA, wild-type mRNA and SNP mRNA of wsv151 or wsv226, RNAi was significantly suppressed, showing that the synonymous SNPs of wsv151 and wsv226 played negative roles in host siRNA pathway due to mismatch of siRNA with its target. In insect cells, the mismatch, caused by synonymous SNPs of wsv151 or wsv226, between siRNA and its target inhibited the host RNAi. Furthermore, the data revealed that the co-injection of SNP siRNA and wild-type siRNA of wsv151 or wsv226 into WSSV-infected shrimp led to a significant increase of WSSV copies compared with that of SNP siRNA alone or wild-type siRNA alone, indicating that the synonymous SNPs of viral genes could be a strategy of virus escaping host siRNA pathway in shrimp in vivo.

## Introduction

Single nucleotide polymorphism (SNP) mainly refers to a DNA or RNA sequence polymorphism caused by the variation of a single nucleotide (1). Generally, SNPs are substitutions, insertions, or deletions of individual bases in the genetic sequence or in non-coding sequences of a DNA. According to the locations, SNPs can be divided into coding SNP and non-coding SNP (2). Based on the change of amino acid or not, the coding SNPs consist of synonymous SNP and non-synonymous SNP (2). The synonymous SNP means that the changed base of a coding sequence does not change its coding protein sequence. At present, SNPs have been widely used as genetic markers to distinguish species, to assess sex ratio of fish populations and to calculate the genomic breeding value in aquaculture (1-4). Recently, it is found that the coding SNPs can influence the health of human being (5, 6). In hepatitis C, the IFNL3 (rs4803217) SNP affects virus infection (6). Many promoter-related SNPs have a relationship with human health (7, 8). As reported, the microRNA (miRNA) pathway can also be influenced by the miRNA-related SNPs (9, 10). The 3’ untranslation region (3’UTR) SNP (rs1048638) of carbonic anhydrase IX gene, the target of miR-34a, can promote hepatocellular carcinoma (9). The bipolar disorder is influenced by SNP rs3749034 which is associated with the hsa-miR-504 pathway (10). In contrast to non-synonymous SNPs, the synonymous SNPs have not been extensively characterized. Although some investigations reveal that synonymous SNPs are associated with diseases (8, 9, 11), the mechanism of synonymous SNPs in diseases remains unclear.

It has been found that a base mutation of a siRNA has a great effect on antiviral RNAi (RNA interference) response of host by impairing the base pair (12, 13), implying that synonymous SNPs may play important roles in the siRNA pathway. RNAi is a vital strategy of animal immune responses to viruses and other foreign genetic materials, especially in invertebrates which lack adaptive immunity (12-18). In shrimp, the vp28-siRNA, which targets the vp28 gene of white spot syndrome virus (WSSV), is capable of mediating sequence-specific gene silencing (12). However, a base mutation of vp28-siRNA suppresses the silencing capacity of vp28-siRNA. It is found that the target recognition by siRNA proceeds via 5’ to 3’ base-pairing propagation with proofreading (14). A base change in the 5’ seed region of a siRNA affects the initial target recognition, while a base change in the 3’ supplementary region of a siRNA promotes the dissociation of RISC (RNA induced silencing complex) before target cleavage (14). These findings suggest that synonymous SNPs can take great effects on the siRNA-mediated RNAi pathway. RNAi, an evolutionarily conserved mechanism in eukaryotes, functions in the antiviral defense of animals through the cleavage and degradation of target viral mRNAs (15-18). In this context, synonymous SNPs may affect the antiviral siRNA pathway of animals. However, this issue is not addressed at present.

To explore the role of synonymous SNPs in antiviral RNAi response, the synonymous SNPs of WSSV were characterized in shrimp in the present study. The results indicated that the synonymous SNPs could be employed by virus to suppress the host’s antiviral RNAi.

## Materials and methods

### Shrimp culture and WSSV challenge

Shrimp (*Marsupenaeus japonicus*), 10 to 12 cm in length, were cultured in groups of 20 individuals in a tank filled with seawater at 25°C. To make sure that shrimp were virus-free before experiments, PCR was conducted to detect WSSV in shrimp using WSSV-specific primers (5’-TATTGTCTCTCCTGACGTAC-3’ and 5’-CACAT TCTTCACGAGTCTAC-3’). DNA was extracted from shrimp with the SQ tissue DNA kit (Omega-Bio-Tek, USA) according to the manufacturer’s instructions. The virus-free shrimp were infected with WSSV (10^5^ copies/ml) by injection (100 μl WSSV inoculum/shrimp) into the lateral area of the fourth abdominal segment. The WSSV isolate used in this study was the Chinese mainland strain (WSSV-CN, GenBank accession no. AF332093.1). At different times post-infection (0, 2, 4, 6, 12, 24, 36, 48, and 72 h), hemocytes of three shrimp, randomly collected for each treatment, were collected for later use.

### Isolation of cytoplasm from shrimp hemocytes

Shrimp homocytes were collected. After centrifugation at 1000×g (4°C) for 10 min, the homocytes were incubated with lysis buffer [10 mM HEPES (2-[4-(2-hydroxyethyl)-1-piperazinyl] ethane sulfonic acid)-KOH, 10 mM MgCl_2,_ 10 mM KCl, 1 mM DTT (dithiothreitol), pH 7.9] for 10 min at 4°C. Subsequently the lysate was centrifuged at 4000×g for 10 min at 4°C. The supernatant (cytoplasm) was collected.

### Western blot analysis

The proteins were separated by 15% SDS-polyacrylamide gelelectrophoresis and then transferred onto a nitrocellulose membrane. The membrane was blocked with 5% nonfat milk in Tris-buffered saline (TBST) (10 mM Tris-HCl, 150 mM NaCl, 20% Tween 20, pH7.5) for 2 h at room temperature, followed by incubation with a primary antibody overnight. After washes with TBST, the membrane was incubated with horseradish peroxidase-conjugated secondary antibody (Bio-Rad, USA) for 2 h at room temperature. The membrane was detected by using a Western Lightning Plus-ECL kit (Perkin Elmer, USA). The antibody against Histone 3 was purchased from Beyotime Biotechnology (China).

### RNA extraction and complementary DNA (cDNA) synthesis

Total RNAs were extracted using the mirVana™ RNA isolation kit according to the manufacturer’s instructions (Ambion, Foster City, USA). To exclude any DNA contamination, the RNAs were treated with RNase-free DNase I (Takara, Shiga, Japan) at 37°C for 30 min. First-strand cDNA synthesis was performed using total RNAs according to the manufacturer’s guidelines for the PrimeScript™1st strand cDNA Synthesis kit (Takara, Shiga, Japan).

### Detection of U6 with PCR

PCR was conducted to detect U6 using U6-specific primers (5’-GGTCTCACTG ACTGTGAC-3’ and 5’-AGTAGCAGTCT CACGAGTCTAC-3’). PCR condition was 94°C for 5 min, followed by 25 cycles of 94°C for 30 s, 54°C for 40 s and 72°C for 1 min, with a final elongation at 72°C for 10 min.

### Analysis of WSSV mRNA SNPs

To obtain a global insight into the characteristics of WSSV mRNA SNPs, total RNAs were extracted from cytoplasm of WSSV-infected shrimp hemocytes. Subsequently RNAs were sequenced by Biomarker Technologies (Beijing, China) with a GA-I genome analyser (Illumina, San Diego, CA, USA). RNA assembly was conducted using TRINITY software (Biomarker Technologies, China). After assembly of RNA-seq data, 7.7 GB raw data was processed to remove the sequences of adapter, ploy-N and low-quality reads. Then the clean reads were analyzed to discover SNPs with the reference sequence of WSSV (GenBank accession no. AF332093.1). The prediction of small interference RNAs (siRNAs) was conducted (https://rnaidesigner.lifetechnologies.com/rnaiexpress/sort.do).

### Northern blot

After separation of small RNAs on a denaturing 15% polyacrylamide gel containing 7 M urea or mRNAs on a 1% agarose gel, RNAs were transferred to a Hybond N+ nylon membrane (Amersham Biosciences) for 1 h at 400 mA, followed by ultraviolet cross-linking. The membrane was prehybridized in DIG (digoxigenin) Easy Hyb granule buffer (Roche, Basel, Switzerland) for 0.5 h at 42 °C and then hybridized with DIG-labeled wsv151 siRNA probe (5’-TCAGCAAGTATGCAGTGT C-3’), wsv226 siRNA probe (5’-GCTACACACAGTATTGTAC-3’), wsv151 mRNA probe (5’-TTCACTTTCAATCTTGGTTCTGC-3’), wsv226 mRNA probe (5’-TCGC TTCAATACCAGATCGTGGAC-3’) or U6 probe (5’-GGGCCATGCTAATCTTCTC TGTATCGTT-3’) at 42 °C overnight. Subsequently the detection was performed with the DIG High Prime DNA labeling and detection starter kit II (Roche).

### SiRNA-mediated cleavage of mRNA *in vitro*

Total RNAs were extracted from shrimp lymphoid organs or hemocytes at 48 h after WSSV infection using a mirVanaTM miRNA isolation kit (Ambion, USA) according to the manufacturer’s instructions. To remove any genomic DNA contamination, total RNA extracts were treated with RNase-free DNase I (Takara, Shiga, Japan) at 37°C for 30 min. First-strand cDNA synthesis was performed using total RNAs according to the manufacturer’s guidelines for PrimeScript 1st strand cDNA Synthesis Kit (Takara, Japan). The first-strand cDNA was used as a template for PCR amplification of wsv151 or wsv226 with sequence-specific primers (wsv151, 5’-GAGGAAGAAAATGTCTACCCCAAA-3’ and 5’-CTGCGTCTCAATTGAAGTGATGTAAAAAT-3’; wsv226, 5’-GTGCGTTTTCAAAGTATAAGAAA-3’ and 5’-CGAGAGAATAAGAATAAAGTGTAT-3’). The SNPs of wsv151 and wsv226 were obtained by PCR using sequence-specific primers (wsv151, 5’-TATGCAGTGACAA TGAGTATCAAAAA-3’ and 5’-ACTCATTGTCACTGCATACTTGCCTC-3’; wsv226, 5’-TAAGCTACATACAGTATTGTACGATAAAG-3’ and 5’-CAATACTGTA TGTAGCTTACATTTAGGTATTA-3’). The PCR products were cloned into pEASY-Blunt Simple vector (TransGen Biotech, China). Then mRNAs were synthesized *in vitro* using a commercial T7 kit according to the manufacturer’s instructions (TaKaRa, Japan).

SiRNAs used were wsv151-siRNA (5’-CGUUCAUACGUCACAGUUA-3’), wsv151-SNP-siRNA (5’-CGUUCAUACGUCACUGUUA-3’), wsv226-siRNA (5’-C GAUGUGUGUCAUAACAUG-3’) and wsv226-SNP-siRNA (5’-CGAUGUAUGU CAUAACAUG-3’). The siRNAs were synthesized using T7 kit according to the manufacturer’s manual (TaKaRa, Japan).

To conduct the siRNA-mediated cleavage of mRNA, the shrimp Ago2 complex was prepared by immunoprecipitation using Ago2-specific antibody in lysis buffer (30 mM HEPES-KOH, pH 7.4), followed by incubation with protein A (Bio-Rad, USA) at 4 °C overnight. The isolated Ago2 complex was dissolved in 500 μl reaction buffer [100 mM KOAc, 2 mM Mg(OAc)_2_] containing 1 mM DTT and protease inhibitor cocktail (Roche, Switzerland) at 4 °C, followed by the addition of 12.5 μl 40×reaction mix [20 μl 500 mM creatine monophosphate, 20 μl amino acid stock (Sigma,USA), 2 μl 1M DTT, 2 μl 20 U/μl RNasin (Promega, USA), 4 μl 100 mM ATP (TaKaRa, Japan), 1 μl 100 mM GTP (TaKaRa), 6 μl 2 U/μl creatine phosphokinase (Cal-Biochem, Germany)]. Then 50 nM siRNA, 100 nM mRNA and the isolated Ago2 complex were incubated at 37 °C for different time (0, 5, 10 and 15min). The reaction was stopped using proteinase K (Shanghai Generay Biotech Co, Ltd. China) by incubation at 37 °C for 5 min. The mRNAs were separated on 1% agarose gel.

### Cell culture, transfection and fluorescence assays

Insect High Five cells (Invitrogen, USA) were cultured in Express Five serum-free medium (Invitrogen) containing L -glutamine (Invitrogen) at 27 °C. At 70% confluence, the insect cells were co-transfected with wild-type siRNA (100 pM) or SNP siRNA (100 pM) and EGFP-wild-type-wsv151/wsv226 (6 μg/ml) or EGFP-SNP-wsv151/wsv226 (6 μg/ml). The constructs were generated by cloning wsv151 and wsv226 mRNAs into the pIZ/V5-His vector (Invitrogen). The recombinant plasmids were confirmed by sequencing. All transfections were carried out in triplicate with Cellfectin transfection reagent (Invitrogen) according to the manufacturer’s protocol. At 36h after co-transfection, the fluorescence intensity of cells was evaluated with a Flex Station II microplate reader (Molecular Devices, USA) at 490/510 nm excitation/emission (Ex/Em). The experiments were biologically repeated three times.

### RNAi assay in shrimp

The small interfering RNA (siRNA) specifically targeting wsv151 (wsv151-siRNA, 5’-CGUUCAUACGUCACAGUUA-3’) or wsv226 (wsv226-siRNA, 5’-CG AUGUGUGUCAUAACAUG-3’) was used in the RNAi assay in shrimp. The synonymous SNP siRNAs (wsv151-SNP-siRNA, 5’-CGUUCAUACGUCACUGUUA -3’; wsv226-SNP-siRNA, 5’-CGAUGUAUGUCAUAACAUG-3’) were included in the RNAi assay. The siRNAs were synthesized using a T7 kit according to the manufacturer’s manual (TaKaRa, Japan). The synthesized siRNAs were quantified by spectrophotometry. Shrimp were co-injected with WSSV (10^4^ copies/shrimp) and siRNA (4nM). At different time after injection, three shrimp were randomly selected from each treatment. The shrimp hemocytes from each treatment were collected for later use.

### Quantification of WSSV copies

Quantitative real-time PCR was used to examine the WSSV copies in shrimp. The genomic DNA of WSSV was extracted with a SQ tissue DNA kit (Omega Bio-tek, Norcross, GA, USA) according to the manufacturer’s instruction. The extracted DNA was analyzed by quantitative real-time PCR with WSSV-specific primers and WSSV-specific TaqMan probe (5’-FAM-TGCTGCCGTCTCCAA-TAMRA-3’) as described previously (19). The linearized plasmid containing a 1400-bp DNA fragment from the WSSV genome was used as the internal standard of quantitative real-time PCR (19). The PCR procedure was 95°C for 1 min, followed by 40 cycles of 95°C for 30 s, 52°C of 30 s, and 72°C for 30 s.

### Statistical analysis

To calculate the mean and standard deviation, the numerical data from three independent experiments were analyzed by one-way analysis of variance (ANOVA). The differences between the different treatments were analyzed by t-test.

## Results

### Existence of SNP in WSSV

To explore the role of SNP in virus infection, the SNPs in cytoplasm of WSSV-infected shrimp hemocytes were characterized. Western blot analysis showed that the isolated cytoplasm of WSSV-infected shrimp hemocytes had no contamination of nucleus (Fig 1A). Then the extracted RNAs from the isolated cytoplasm, without contamination of nuclear RNAs (Fig 1B), were subjected to RNA-seq analysis.

**Fig 1.**
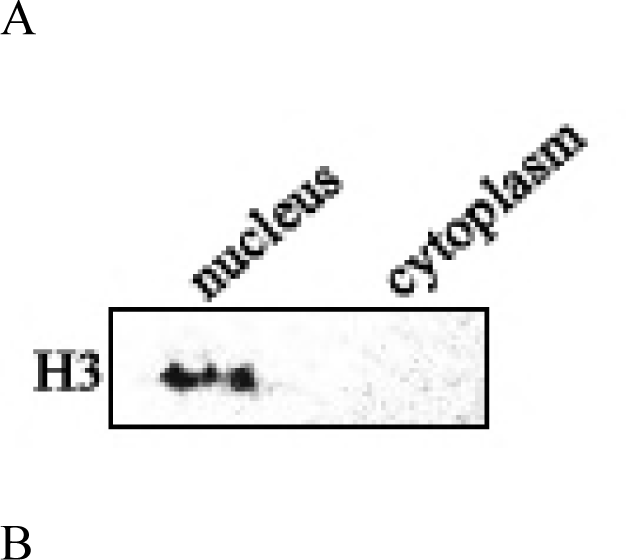
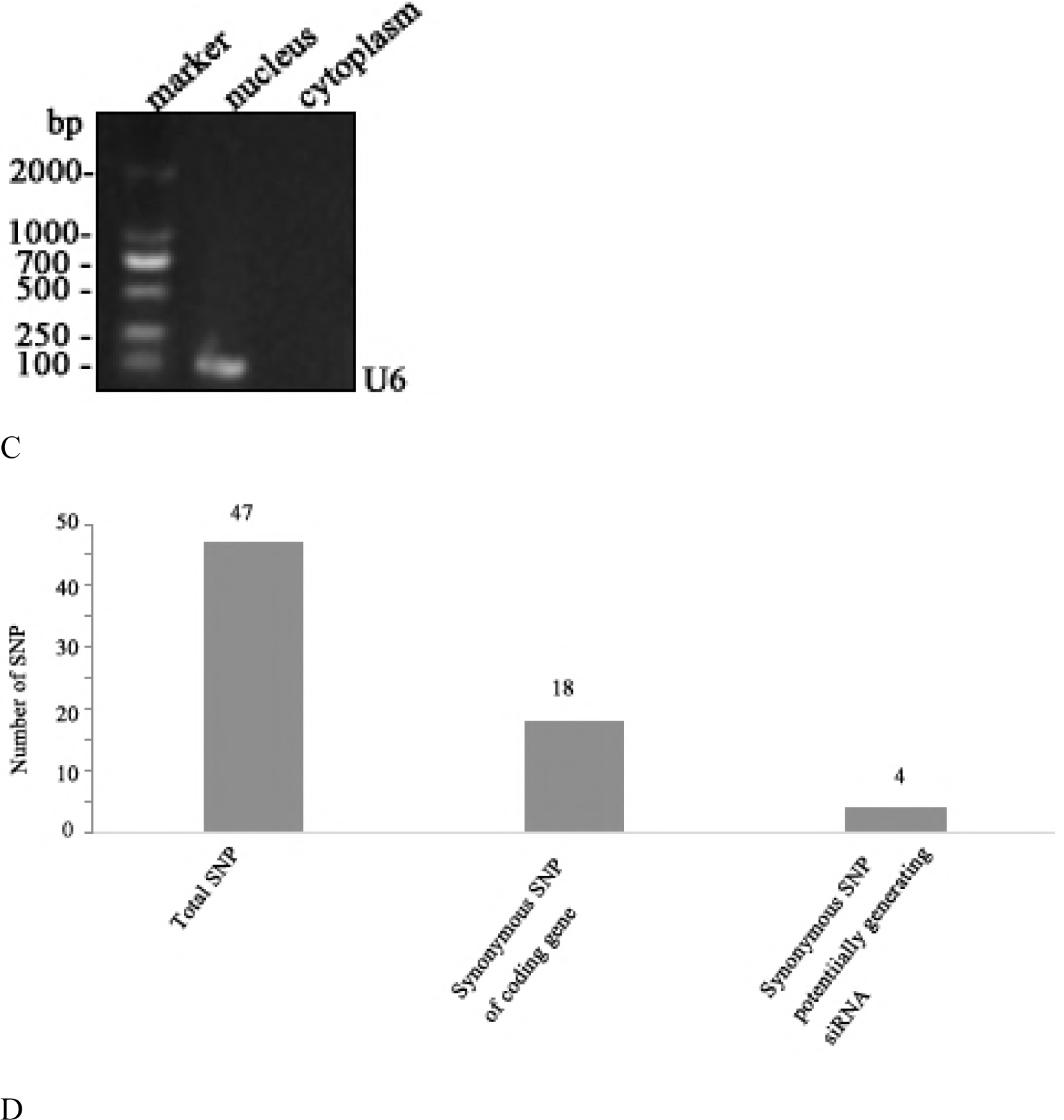
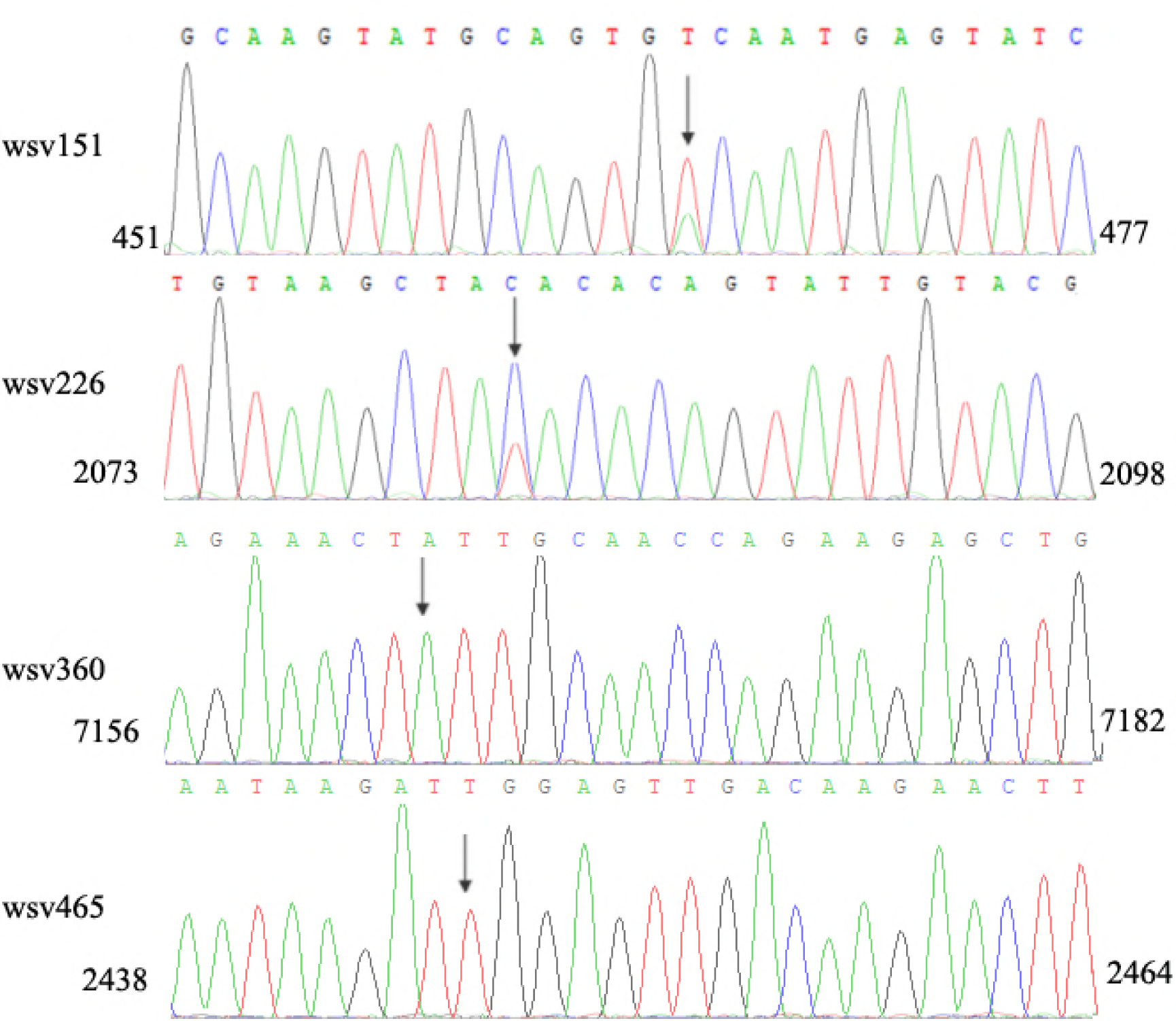
Existence of SNP in WSSV. (A) Western blot analysis of the isolated cytoplasm. Shrimp were infected with WSSV. At 48h post-infection, the shrimp hemocytes were collected and subjected to the isolation of cytoplasm. To exclude the contamination of nucleus, the isolated cytoplasm was detected using Western blot with anti-histone 3 (H3) IgG. (B) PCR detection of RNAs extracted from the isolated cytoplasm RNA. RNAs were extracted from the isolated cytoplasm and then reversely transcribed into cDNAs, followed by PCR using U6-specific primers. The PCR products were analyzed using agarose gel electrophoresis. (C) The number of synonymous SNPs. (D) Identification of synonymous SNPs of WSSV mRNAs. The RNAs extracted from hemocytes of WSSV-infected shrimp were sequenced. The arrows indicated the SNP sites and the numbers showed the positions of the coding sequences of viral genes.

The sequence analysis showed that all the 180 open reading frames (ORFs) of WSSV were found in the RNA-seq analysis. Among them, 16 WSSV mRNAs possessed 47 SNPs (Fig 1C). Of the 47 SNPs, there were 18 synonymous SNPs from 10 genes (Fig 1C). As well known, synonymous SNPs of an mRNA could not take effects on its protein-coding sequence. The synonymous SNPs of a viral mRNA might function in antiviral siRNA pathway to escape the targeting by siRNA. The sequence analysis indicated that 4 WSSV mRNAs (wsv465, wsv360, wsv226 and wsv151) containing 4 synonymous SNPs were predicted to potentially generate siRNAs employed by shrimp (Fig 1C).

To investigate the existence of the 4 synonymous SNPs, the RNAs extracted from hemocytes of WSSV-infected shrimp were sequenced. The sequence analysis showed that two SNPs (wsv151 and wsv226) existed in WSSV-infected shrimp (Fig 1D). However, the wsv360 SNP and the wsv465 SNP were not detected (Fig 1D). The sequencing of WSSV genomic DNA generated the same results, confirming the existence of SNPs of wsv151 and wsv226.

Taken together, these findings revelaed that SNP existed in WSSV and that 2 synonymous SNPs (wsv151 and wsv226) might function in the siRNA pathway of shrimp.

### Influence of viral synonymous SNPs on host siRNA pathway

In an attempt to reveal the roles of viral synonymous SNPs in host siRNA pathway, the wsv151 siRNA and wsv226 siRNA of WSSV were examined in hemocytes of WSSV-infected shrimp. Northern blot results indicated that shrimp could generate the wsv151 siRNA and wsv226 siRNA during WSSV infection (Fig 2A). To explore the influence of synonymous SNPs of wsv151 and wsv226 mRNAs on shrimp siRNA pathway, the sequences of wild-type siRNAs and mRNAs of wsv151 and wsv226 were mutated, generating SNP-siRNAs and SNP-mRNAs which contained the synonymous SNPs of wsv151 and wsv226, respectively. Then the synthesized wild-type siRNA (WT siRNA) or SNP siRNA and WT mRNA or SNP mRNA were incubated with the isolated shrimp Ago2 complex, followed by separation of mRNAs on agarose gel. The results indicated that wild-type siRNA and SNP-siRNA of wsv226 could mediate the cleavage of wild-type mRNA and SNP-mRNA of wsv226, respectively (Fig 2B). However, the cleavage of wsv226 wild-type mRNA mediated by wsv226 SNP-siRNA and the cleavage of wsv226 SNP-mRNA mediated by wsv226 wild-type siRNA were inhibited (Fig 2B). For wsv151, the viral synonymous SNP yielded the similar results (Fig 2C). These data revealed that the viral synonymous SNPs suppressed the host siRNA pathway.

**Fig 2.**
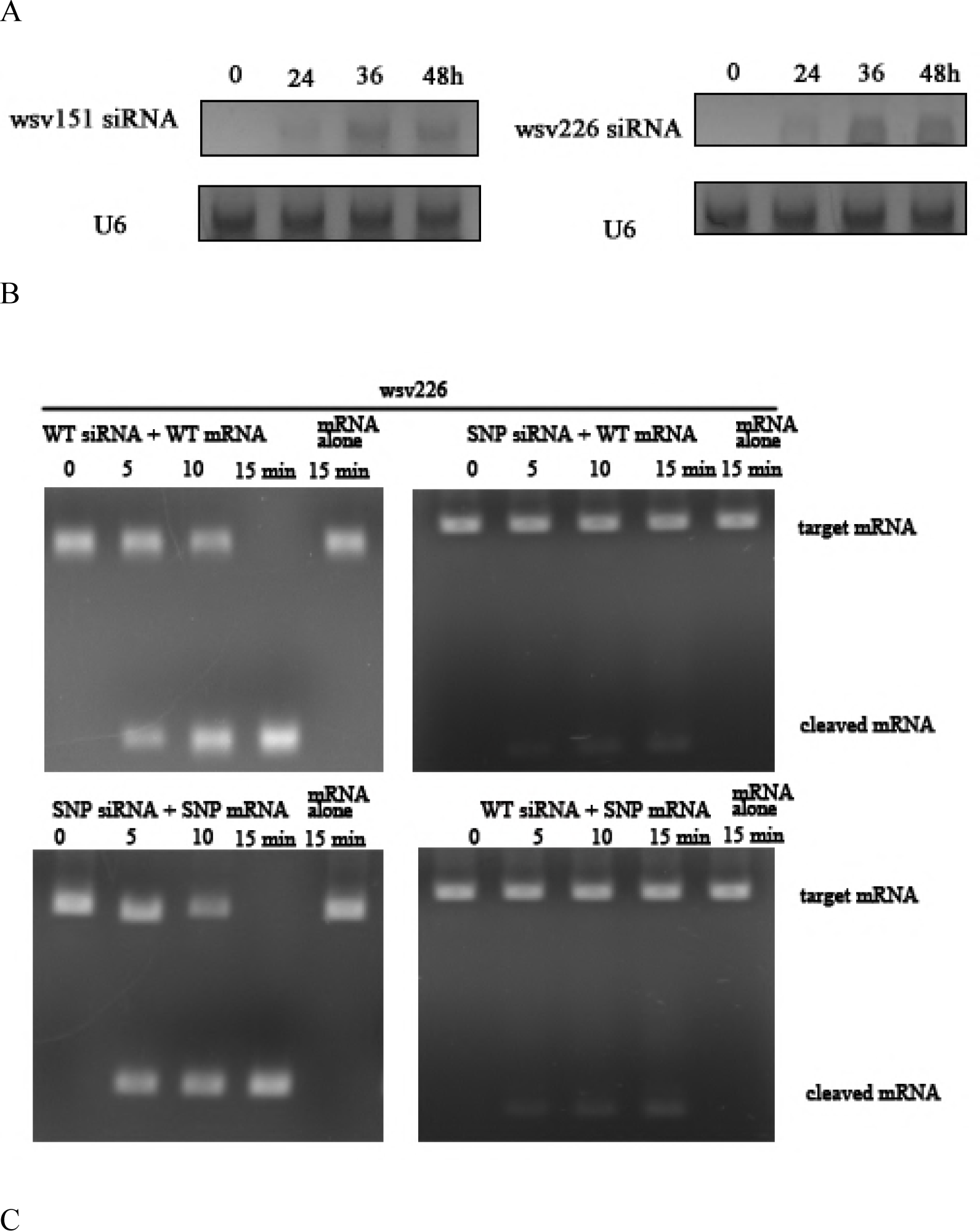
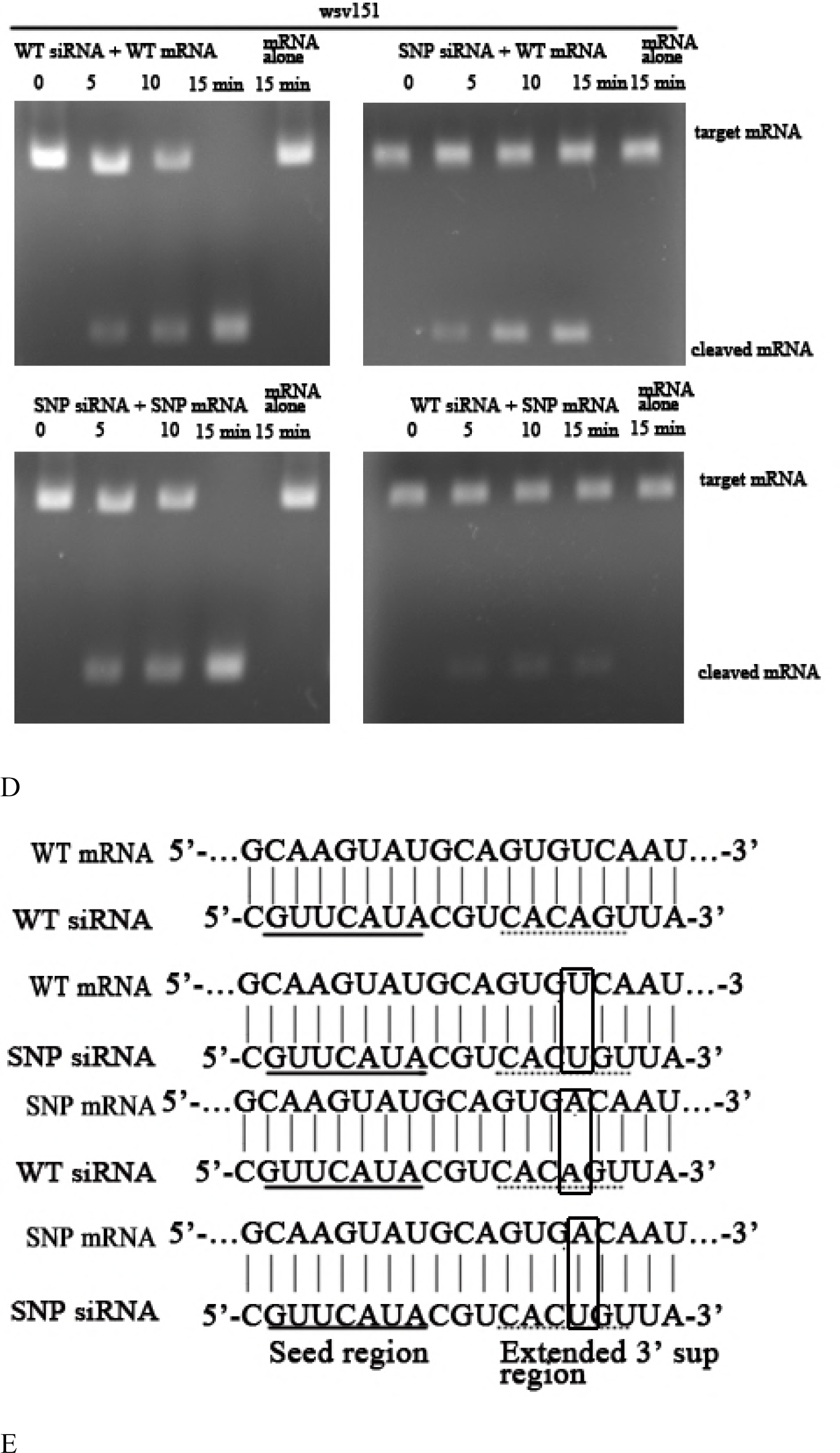
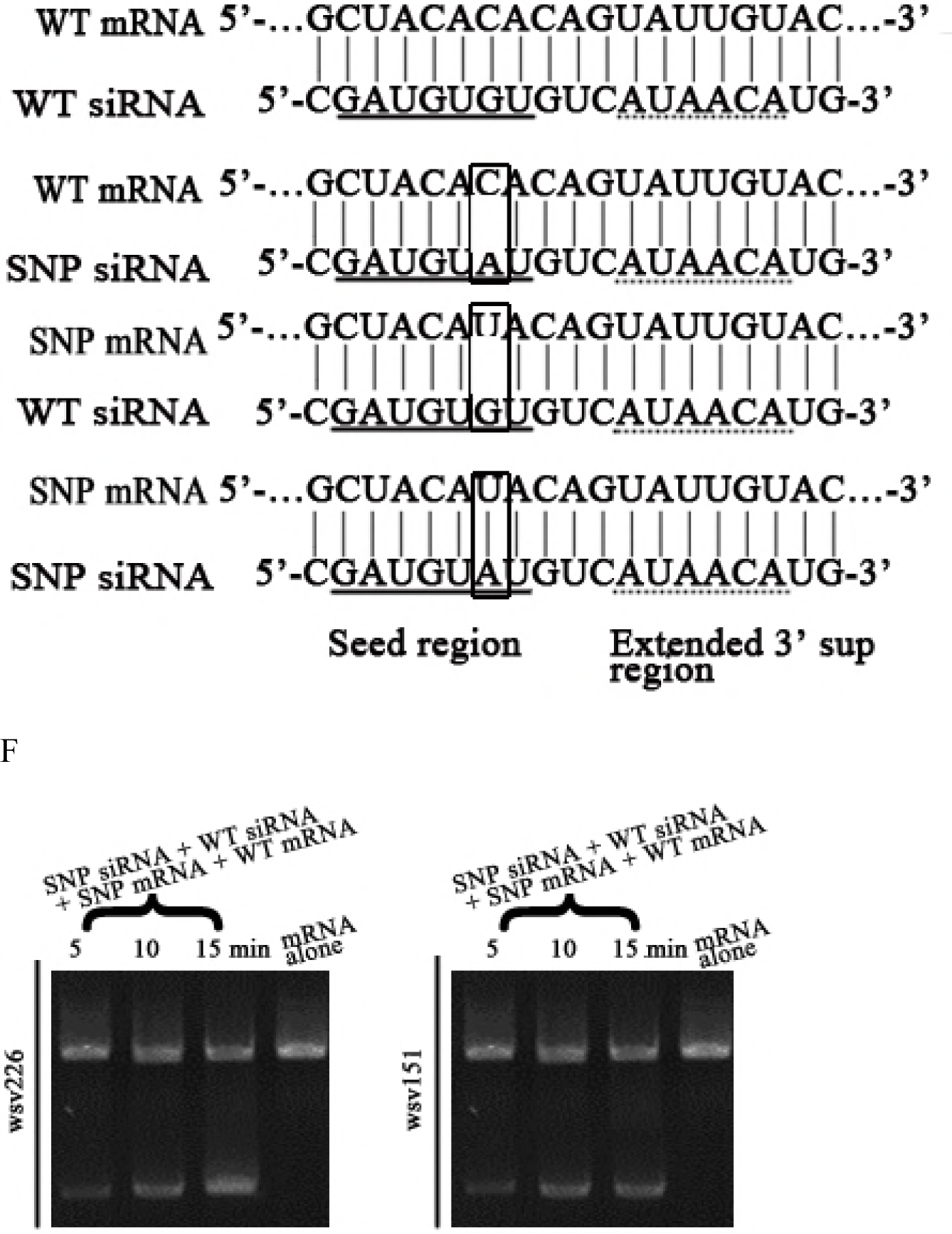
Influence of viral synonymous SNPs on host siRNA pathway. (A) The detection of siRNA in WSSV-infected shrimp. Shrimp were infected with WSSV. At different times post-infection, wsv151 siRNA and wsv226 siRNA were detected using hemocytes of WSSV-infected shrimp by Northern blotting. U6 was used as a control. (B) The influence of synonymous SNP of wsv226 mRNA on shrimp siRNA pathway. The wild-type siRNA (WT siRNA) or SNP siRNA and wild-type mRNA (WT mRNA) or SNP mRNA were incubated with the isolated shrimp Ago2 complex. At different times after incubation, the mRNA was separated by agarose gel electrophoresis. Wsv226 mRNA alone was used as a control. (C) The role of synonymous SNP of wsv151 mRNA in shrimp siRNA pathway. The WT siRNA or SNP siRNA, WT mRNA or SNP mRNA and the isolated shrimp Ago2 complex were incubated for different time. The mRNA cleavage was examined using agarose gel electrophoresis. Wsv151 mRNA alone was used as a control. (D) The sequence analysis of wild-type and SNP mRNA and siRNA of wsv151. Boxes indicated the SNPs. The seed region and extended 3’ sup (supplementary) region of siRNA were underlined with solid and dashed lines, respectively. (E) The sequence analysis of wild-type and SNP mRNA and siRNA of wsv226. (F) The impact of synonymous SNPs on RNAi in the presence of SNP mRNA, SNP siRNA, WT mRNA and WT siRNA. SNP mRNA, SNP siRNA, wild-type (WT) mRNA and WT siRNA were mixed and then incubated with the isolated shrimp Ago2 complex. At different times after incubation, the mRNA was separated by agarose gel electrophoresis. As a control, mRNA alone was included in the assays.

To reveal the locations of synonymous SNPs of wsv151 and wsv226, the sequences of wild-type and SNP mRNAs and siRNAs were compared. The results showed that the synonymous SNP of wsv151 was located in the extended 3’ supplementary region of siRNA (Fig 2D). For the wsv226 synonymous SNP, it was located in the seed region of siRNA (Fig 2E). As reported, the seed region (positions 2-8) and the extended 3’ supplementary region (positions 12-17) of a siRNA were essential for the siRNA-mediated gene silencing (14). Therefore, the synonymous SNPs of wsv151 and wsv226 could inhibit the host siRNA pathway.

The sequence showed that the ratio of wild-type mRNA to SNP mRNA of wsv151 was 8 to 5, while the ratio of wild-type mRNA to SNP mRNA of wsv226 was 4 to 7. These data indicated the co-existence of wild-type mRNAs and SNP mRNAs of WSSV in natural conditions. Based on prediction, it was found that SNP siRNA could be produced from SNP mRNA of wsv151 or wsv226, showing the co-existence of wild-type siRNA and SNP siRNA. Therefore, wild-type mRNA, SNP mRNA, wild-type siRNA and SNP siRNA of wsv151 or wsv226 simultaneously existed in WSSV-infected shrimp. The results showed that RNAi was significantly suppressed in the presence of SNP siRNA, wild-type siRNA, wild-type mRNA and SNP mRNA (Fig 2F).

The above findings indicated that the synonymous SNPs of viral mRNA played negative roles in host siRNA pathway.

### Suppression of siRNA pathway by viral synonymous SNP in insect cells

To explore the effects of synonymous SNPs of wsv151 mRNA and wsv226 mRNA on siRNA pathway in cells, the insect High Five cells were co-transfected with wild-type siRNA (WT siRNA) or SNP-siRNA and EGFP-wild-type-wsv151 mRNA, EGFP-SNP-wsv151 mRNA, EGFP-wild-type-wsv226 mRNA or EGFP-SNP-wsv226 mRNA. At 36 h after co-transfection, the wsv151 mRNA, the wsv226 mRNA and the fluorescence intensity of insect cells were evaluated.

The results of Northern blot showed that when the insect cells were co-transfected with WT siRNA and WT mRNA of wsv226 or SNP siRNA and SNP mRNA of wsv226, the wsv226 mRNA level was significantly decreased (Fig 3A). However, the transfection of insect cells with WT siRNA and SNP mRNA of wsv226 or SNP siRNA and WT mRNA of wsv226 did not change the wsv226 mRNA level (Fig 3A). In the simultaneous existence of WT siRNA, SNP siRNA, WT mRNA and SNP mRNA of wsv226, the siRNA pathway was significantly inhibited compared with the control (Fig 3A). The fluorescence intensity of insect cells generated the similar results (Fig 3B). These data indicated that the viral synonymous SNP of wsv226 escaped the siRNA pathway in insect cells.

**Fig 3.**
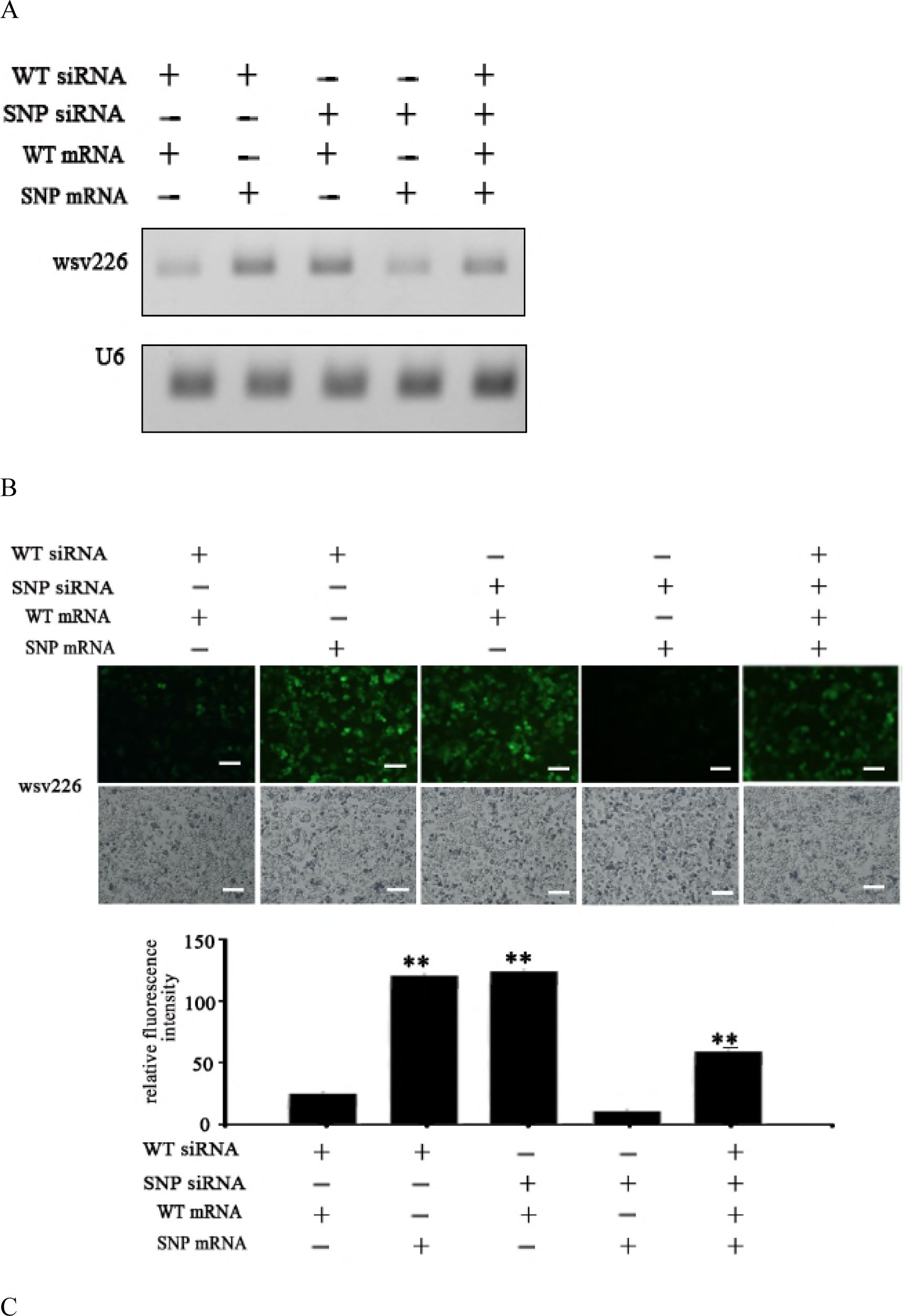
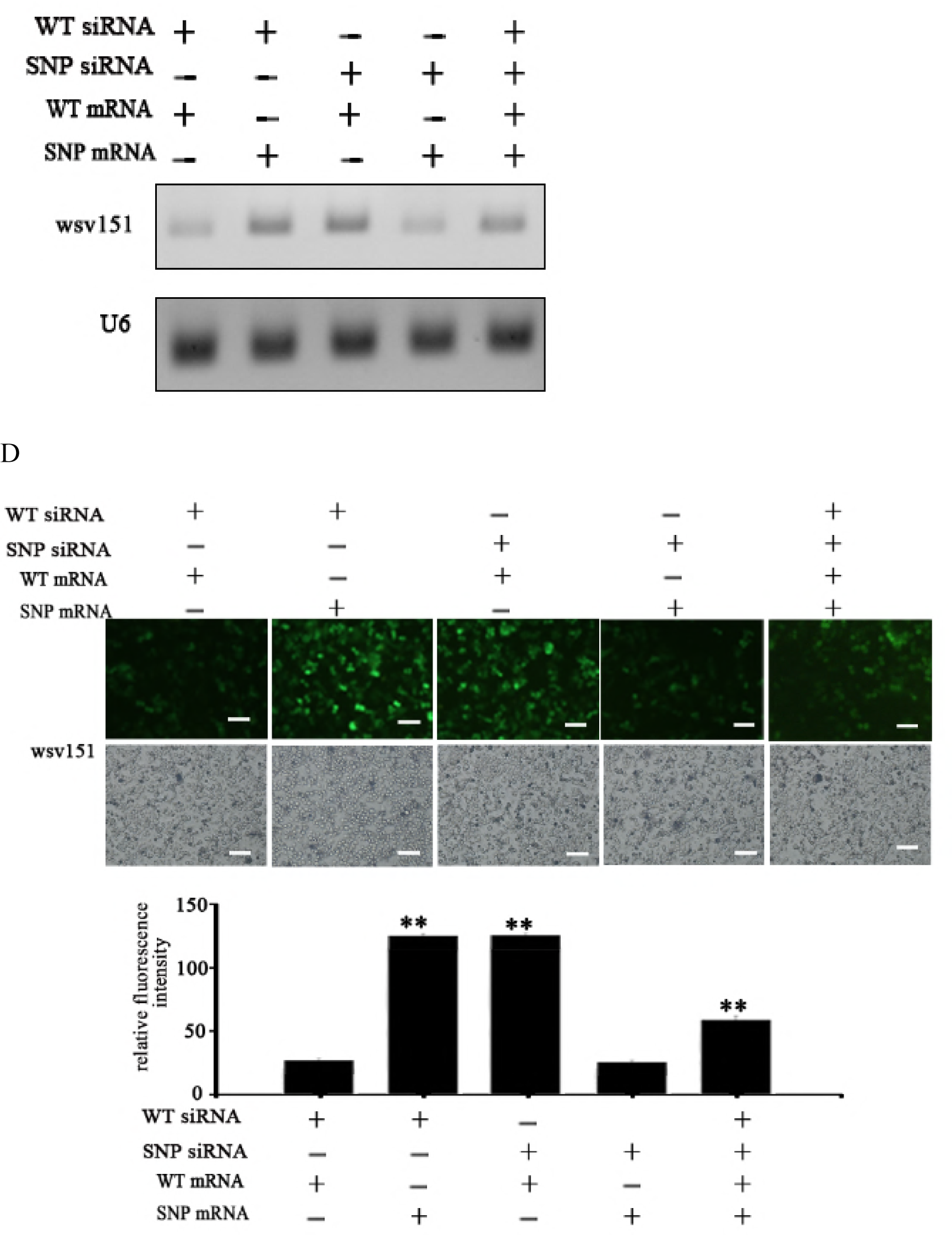
Suppression of siRNA pathway by viral synonymous SNP in insect cells. (A) Influence of synonymous SNP of wsv226 mRNA on siRNA pathway in insect cells. The insect High Five cells were co-transfected with wild-type siRNA (WT siRNA), SNP siRNA, EGFP-WT-mRNA and/or EGFP-SNP-mRNA of wsv226. At 36 h after co-transfection, the wsv226 mRNA level was examined by Northern blot. U6 was used as a control. Probes were indicated on the left. (B) Evaluation of viral synonymous SNP of wsv226 mRNA on siRNA pathway in insect cells by examination of fluorescence intensity of insect cells. The treatments were indicated on the top. Scale bar, 50 μm. (C) Role of synonymous SNP of wsv151 mRNA in siRNA pathway in insect cells. At 36 h after co-transfection of insect cells with wild-type (WT) siRNA, SNP siRNA, EGFP-WT-mRNA and/or EGFP-SNP-mRNA of wsv151, the wsv151 mRNA level was determined by Northern blot. U6 was used as a control. Probes were indicated on the left. (D) Impact of viral synonymous SNP of wsv151 mRNA on siRNA pathway in insect cells by examination of fluorescence intensity of insect cells. The treatments were indicated on the top. Scale bar, 50 μm.

For wsv151, Northern blots indicated that the wsv151 mRNA level was significantly reduced when the insect cells were co-transfected with WT siRNA and WT mRNA of wsv151 or SNP siRNA and SNP mRNA of wsv151 (Fig 3C). However, the co-existence of WT siRNA, SNP siRNA, WT mRNA and SNP mRNA of wsv151, resulted in a significant suppression of RNAi in insect cells (Fig 3A). The fluorescence intensity of insect cells essentially generated the similar results (Fig 3D). These results revealed that the viral synonymous SNP of wsv151 could inhibit the siRNA pathway.

Taken together, the findings showed that the synonymous SNPs of viral mRNAs could escape the siRNA pathway in cells.

### Inhibition of siRNA pathway by viral synonymous SNPs in shrimp *in vivo*

In order to characterize the influence of viral synonymous SNPs on the host siRNA pathway *in vivo*, the synthesized wild-type (WT) siRNA or/and SNP siRNA of wsv151 or wsv226 and WSSV were co-injected into shrimp and then the wsv151 and wsv226 mRNAs were examined. The results indicated that the mRNA levels of wsv151 and wsv226 were significantly decreased in shrimp treated with WT siRNA or SNP siRNA compared with the control (WSSV alone) (Fig 4A and B). When shrimp were co-injected with WT siRNA and SNP siRNA, the expression profiles of wsv151 and wsv226 in WSSV-infected shrimp were similar to the WT siRNA or SNP siRNA treatment (Fig 4A and B). These findings revealed that SNPs of wsv151 and wsv226 could escape the siRNA pathway of shrimp *in vivo*.

**Fig 4.**
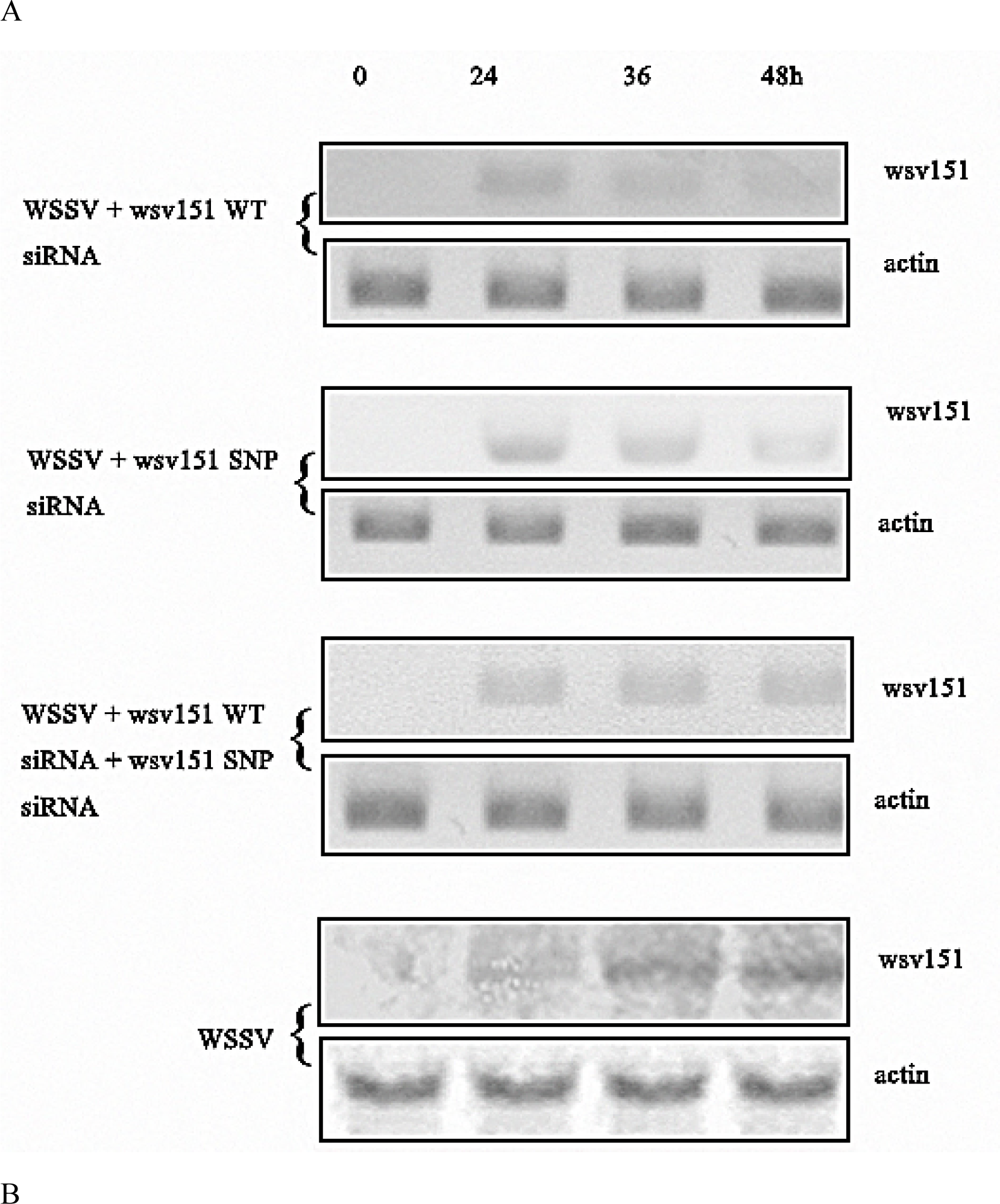
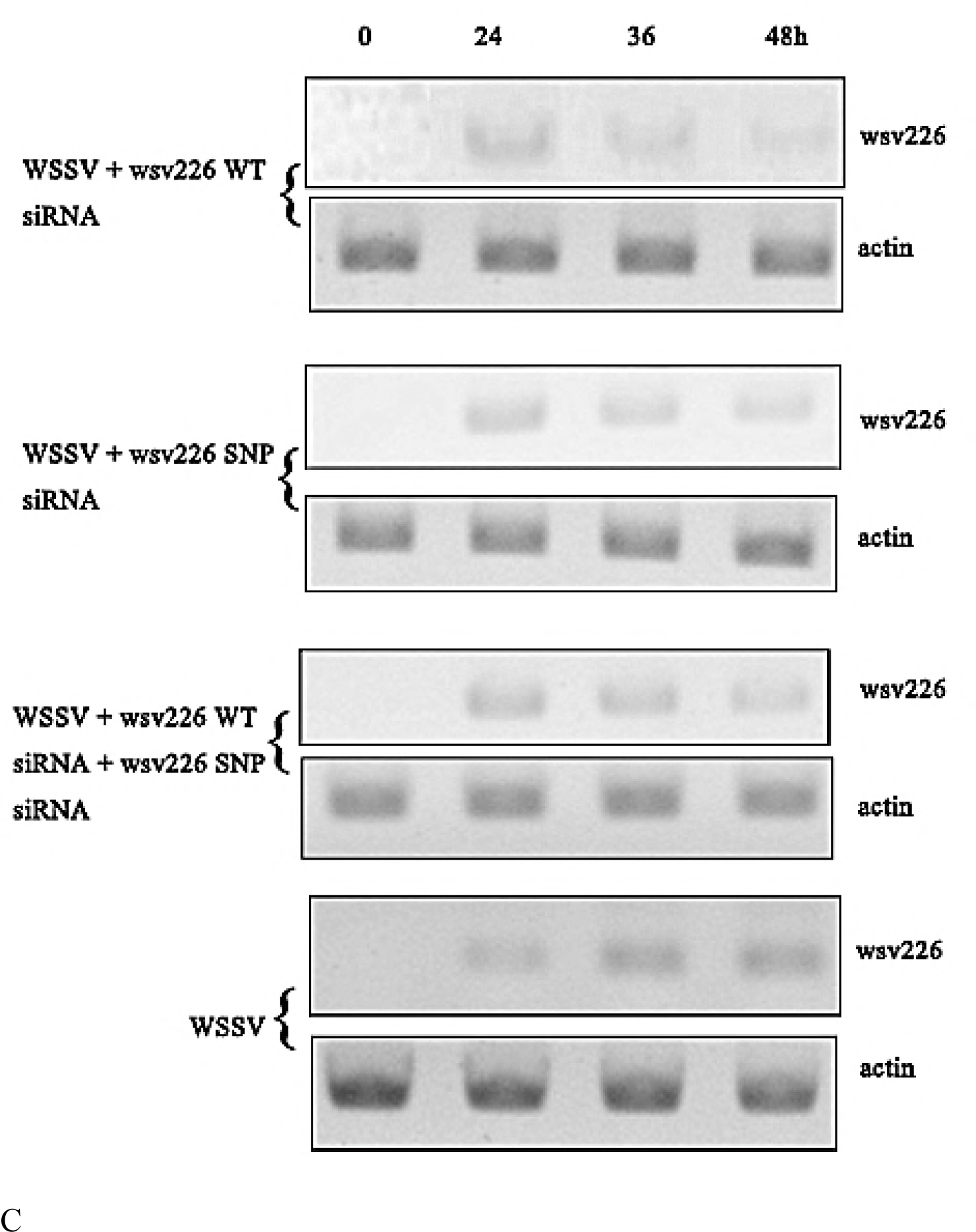
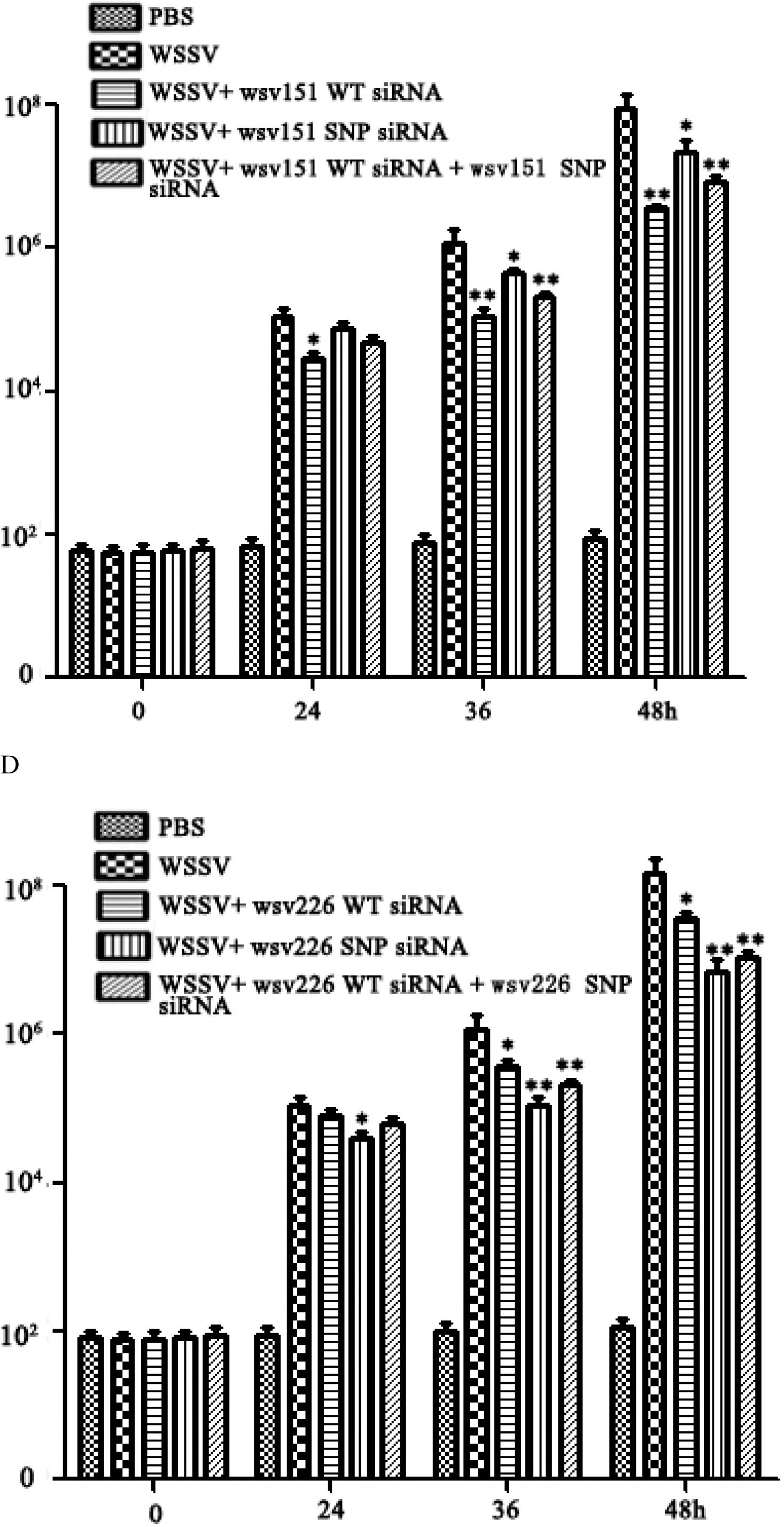
Inhibition of siRNA pathway by viral synonymous SNPs in shrimp *in vivo*. (A) Influence of synonymous SNPs on wsv151 expression in WSSV-infected shrimp. Shrimp were co-injected with WSSV and WT siRNA or/and SNP siRNA of wsv151. WSSV alone was included in the injection as a positive control. At different time after injection, the shrimp hemocytes were analyzed by Northern blotting. β-actin was used as a loading control. Numbers indicated the time post-infection. (B) Impact of synonymous SNPs on the expression of wsv226 in WSSV-infected shrimp. WSSV and siRNA or/and SNP siRNA of wsv226 were injected into shrimp. At different time post-infection, the shrimp hemocytes were subjected to Northern blotting. (C) Detection of WSSV copies in shrimp hemocytes by quantitative real-time PCR. Shrimp were injected with WSSV and wsv151 WT siRNA or/and wsv151 SNP siRNA. WSSV alone and PBS were included in the injection as controls. At different time after injection (0, 24, 36 and 48h), the WSSV copies in shrimp were examined (*, *p*<0.05; **, *p*<0.01). (D) Effects of synonymous SNPs of wsv226 on WSSV infection in shrimp. The treatments were indicated on the top. The numbers showed the time post-infection. Significant differences between treatments were indicated with asterisks (*, *p*<0.05; **, *p*<0.01).

To explore the effects of viral synonymous SNPs on virus infection *in vivo*, shrimp were injected with WSSV and wild-type (WT) siRNA or/and SNP siRNA of wsv151 or wsv226, followed by the detection of WSSV copies. The results showed that the silencing of wsv151 and wsv226 by WT siRNAs led to a significant decrease of WSSV copies compared with the control (WSSV alone) (Fig 4C and D). SNP siRNA of wsv151 or wsv226 yielded the similar results (Fig 4C and D). When WT siRNA and SNP siRNA of wsv151 or wsv226 were co-injected into WSSV-infected shrimp, the WSSV copies were significantly reduced in shrimp compared with the control (WSSV alone) (Fig 4C and D).

The above findings revealed that the synonymous SNPs could be a strategy of virus escaping host siRNA pathway in shrimp *in vivo*.

## Discussion

As reported, SNPs in the non-coding regions of a gene can manifest a higher risk of cancer (20) and can affect mRNA structure and disease susceptibility (21, 22). In the coding region, SNPs may result in a premature stop codon, a nonsense codon in the transcribed mRNA or in a truncated, incomplete and nonfunctional protein product (21). In recent years, it is found that synonymous SNPs are involved in codon usage preference or mRNA splicing (23-25). In the genomes of almost all species, one of the most prominent observations on the non-neutrality of synonymous codons is the correlation between synonymous codon usage bias and the level of gene expression (23). Highly expressed genes tend to have a higher preference toward so-called optimal codons than lowly expressed genes (23). The amounts of cognate tRNAs that bind to optimal codons are significantly higher than the amounts of cognate tRNAs that bind to non-optimal codons in genomes (23). In some cases, synonymous SNPs are associated to mRNA splicing (24-26). Based on their impact on mRNA splicing and protein function, it is suggested that the protein structure features offer an added dimension of information while distinguishing disease-causing and neutral synonymous SNPs (26). In this study, the results showed that the viral synonymous SNPs of wsv1151 and wsv226 played negative roles in shrimp RNAi, indicating the synonymous SNP was employed by virus to escape the host’s RNAi immunity during virus infection. Therefore, our findings contributed a novel function of synonymous SNPs.

It has been demonstrated that the siRNA-mediated RNAi is the major antiviral defense of invertebrates which lack adaptive immunity (27, 28). In the siRNA-induced silencing complex (RISC), the siRNA uses not only its seed region (the 2nd-7th nt) but also its 3’ supplementary region (the 12th-17th nt) (also called extended 3’ supplementary region) (14, 29). The target recognition of siRNA depends on base pairing in the seed region but not in the 3’ supplementary region of the siRNA, showing the importance of seed pairing for initial target recognition (14). It is also found that the mismatch in the 3’ supplementary region of a siRNA prevents RISC from reaching its cleavage-competent conformation and promoting dissociation of RISC without cleavage (14). In the presence of mismatches of siRNA with its target mRNA (wild-type mRNA), the mismatched siRNA and target mRNA can be loaded into RISC. However, the target cleavage does not occur in this RISC (14). In shrimp, the findings show that a single-base mutation in the 6th base of vp28-siRNA leads to the suppression of shrimp antiviral immunity against WSSV (12). In the present study, the results revealed that RNAi was significantly suppressed in the presence of SNP siRNA, wild-type siRNA, wild-type mRNA and SNP mRNA of wsv151 or wsv226. Therefore, in the siRNA-mediated RNAi, a single-base mutation of a siRNA can inactivate the antiviral RNAi immunity. Our *in vitro* and *in vivo* data revealed that WSSV could escape the shrimp RNAi immunity by the single-base mismatch between the seed region or the 3’ supplementary region of a siRNA and its target mRNA during WSSV infection. In this context, our findings presented that the synonymous SNPs played important roles during virus infection, which would be helpful for us to gain insights into the molecular mechanisms in virus-host interactions *in vivo*.

## Acknowledgements

This work was supported by National Natural Science Foundation of China (31430089) and National Program on the Key Basic Research Project (2015CB755903).

